# E-cadherin maintains oral Langerhans cell barrier surveillance to preserve microbiota-dependent immune homeostasis

**DOI:** 10.64898/2026.07.07.736593

**Authors:** Anna Brand, Sarah Angabo, Mariia Antipova, Andressa V. B. Nogueira, Andreas Hiergeist, Philipp Münch, Reem Naamneh, Mai Gara, Matthias Klein, Anna Damanaki, Tobias Bopp, James Deschner, André Gessner, Avi-Hai Hovav, Björn E. Clausen

## Abstract

Langerhans cells (LC) are specialized antigen-presenting cells that form a dense immune surveillance network within the oral epithelium. There, they continuously interact with epithelial cells and the resident microbiota to maintain mucosal homeostasis. A defining feature of LC is their highly dendritic morphology, which enables efficient sampling of the environment at barrier surfaces. Although E-cadherin–mediated adhesion has been implicated in LC–epithelial cell interactions, its role in oral LC biology and periodontal immune homeostasis remains elusive. Here, we investigated the function of E-cadherin on oral LC using CD11c-specific E-cadherin–deficient (CD11c-Ecad^DEL^) mice. Loss of E-cadherin profoundly altered LC morphology throughout the oral mucosa, resulting in reduced dendrite formation and impaired dendrite extension towards the epithelial surface, thereby disrupting interaction with the oral microbiota. While the total number of LC remained unchanged, E-cadherin deficiency significantly altered the relative distribution of LC subsets, characterized by reduced LC1 and increased LC2 populations. E-cadherin–deficiency was associated with pronounced oral dysbiosis, characterized by increased bacterial burden and microbial diversity, as well as a shift away from the commensal-dominated community, particularly through the loss of protective lactobacilli. Transcriptome analysis of gingival tissue revealed inflammatory reprogramming marked by enrichment of NF-κB, TNF, IL-17, Toll-like receptor, and MAPK signaling pathways. Consistently, CD11c-Ecad^DEL^ mice exhibited increased IL-17A production in the gingiva, expansion of αβ and γδ T cells, spontaneous age-dependent alveolar bone loss, and exacerbated inflammatory bone destruction in a model of ligature-induced periodontitis. In summary, our findings reveal that E-cadherin preserves oral LC dendrite organization and microbiota-dependent immune homeostasis, thereby limiting dysbiosis-driven inflammation and periodontal bone loss.

## Introduction

The oral mucosa represents a highly specialized barrier tissue that is continuously exposed to a dense and diverse microbial environment. Maintenance of oral homeostasis therefore requires a delicate balance between immune tolerance towards commensal microorganisms and the ability to mount protective immune responses against pathogens (Hovav 2014, Moutsopoulos and Konkel 2018, Brand, Hovav et al. 2023). Disruption of this balance can result in chronic inflammatory diseases such as periodontitis, a highly prevalent condition characterized by oral dysbiosis, gingival inflammation, and progressive alveolar bone destruction (Hajishengallis 2014, Hajishengallis and Lamont 2021). Increasing evidence indicates that periodontal disease is not driven solely by pathogenic bacteria, but rather emerges from dysregulated host immune responses to a polymicrobial dysbiotic community (Lamont, Koo et al. 2018, Hajishengallis and Lamont 2021).

In this context, antigen-presenting cells such as Langerhans cells (LC) play a critical role in orchestrating immune responses within the gingival epithelium. They account for approximately 2–4% of cells in the oral epithelium and are strategically positioned within the suprabasal epithelial layers, where their characteristic dendritic morphology enables continuous sampling of environmental antigens and interaction with the local microbiota. They act as sentinels capable of recognizing pathogens, internalizing antigens, and migrating to regional lymph nodes (LN) to prime naïve T cells (Capucha, Koren et al. 2018, Hovav 2018, Brand, Hovav et al. 2023). Through these mechanisms, LC contribute not only to antimicrobial immunity but also to immune tolerance towards commensal microbiota. LC in the oral mucosa exhibit functional and developmental characteristics that differ from their counterparts in the skin (Hovav 2018, Brand, Hovav et al. 2023). While epidermal LC are largely derived from embryonic macrophage precursors and maintain themselves through local self-renewal, oral mucosal LC originate from circulating dendritic cell progenitors and monocytes (Merad, Manz et al. 2002, Hoeffel, Wang et al. 2012, Capucha, Mizraji et al. 2015). This dynamic turnover suggests that oral LC are particularly adapted to respond to the constantly changing microbial environment of the oral cavity. On one hand, the local microbiota has a major impact on the development of mucosal LC; on the other hand, LC in turn maintain mucosal homeostasis and prevent tissue destruction (Capucha, Koren et al. 2018). Prolonged ablation of oral LC using Langerin-DTR mice results in an increased total bacterial load, a reduced frequency of regulatory T cells, and exacerbated alveolar bone destruction, demonstrating that LC are crucial for maintaining microbial homeostasis (Capucha, Koren et al. 2018). Like classical dendritic cells, oral LC can be differentiated into CD103^+^ LC type-1 (LC1), CD11b^+^ LC2, and CX3CR1^+^ monocyte-derived LC (moLC) (Capucha, Mizraji et al. 2015). To date, the functional specialization of these subsets in oral mucosal homeostasis and periodontal disease is unknown. The finding that germ-free mice harbor less CD103^+^ LC1 compared to SPF mice, while the number of CD11b^+^ LC2 remains largely unchanged, suggests that primarily CD103^+^ LC1 interact with the oral microbiota (Capucha, Koren et al. 2018).

To perform their immune surveillance function, LC constantly extend their dendrites through the suprabasal epithelial layers towards the surface of the epithelium, enabling efficient antigen capture from the oral environment (Kubo, Nagao et al. 2009, Capucha, Koren et al. 2018). The formation and maintenance of these dynamic dendritic structures require coordinated interactions between LC and the surrounding epithelial cells. One key molecule involved in this process is E-cadherin, a calcium-dependent adhesion molecule expressed by both LC and epithelial cells (Tang, Amagai et al. 1993). Although E-cadherin has long been considered essential for the maintenance of LC within epithelial tissues, we recently demonstrated that the number of epidermal LC and their tissue retention are preserved in the absence of E-cadherin, indicating that E-cadherin is dispensable for LC differentiation and maintenance under steady-state conditions in the skin (Brand, Diener et al. 2020). Interestingly, long-lived epidermal LC require E-cadherin to form their typical dendrites, which could impair their capacity for antigen sampling and immune surveillance of the skin (Kubo, Nagao et al. 2009, Brand, Diener et al. 2020). Whether E-cadherin is required for the maintenance of LC morphology and immune function in the oral epithelium remains unknown. Given the distinct developmental and environmental conditions of mucosal LC compared to epidermal LC, E-cadherin–dependent epithelial interactions may play tissue-specific roles in oral barrier immunity. Understanding how epithelial adhesion molecules such as E-cadherin regulate LC morphology and function in the oral mucosa – both at steady-state and during disease – may provide new insights into the mechanisms governing mucosal immunity and inflammatory responses in periodontal tissues.

## Results

### E-cadherin deficiency alters LC morphology and subset composition in the oral epithelium

To investigate the role of E-cadherin in maintaining oral LC morphology, we performed immunofluorescence staining of epithelial sheets from CD11c-Ecad^DEL^ mice (which lack E-cadherin specifically on all CD11c-expressing cells) and control mice. Oral tissues (gingiva, buccal mucosa, and tongue epithelium) were isolated and stained for MHCII and Langerin to identify LC. In control mice, LC displayed their characteristic dendritic morphology, featuring elongated cell bodies and numerous protruding dendrites, and formed a dense network (Fig. 1a). In contrast, LC in CD11c-Ecad^DEL^ mice exhibited a markedly altered morphology, appearing more rounded with a significant reduction in dendritic extensions (Fig. 1b). This was observed in all oral tissues (Fig. 1, SupplFig. 1). Consistently, histological analysis of tongue cross sections confirmed that LC in CD11c-Ecad^DEL^ mice displayed a less ramified structure and shortened dendrites, and were unable to extend their dendrites to the surface of the epithelium (Fig. 1c). These findings demonstrate that E-cadherin is crucial for maintaining the dendritic morphology of LC in the oral mucosa. To determine whether the loss of dendrites in E-cadherin-deficient LC affects LC homeostasis under steady-state conditions, we analyzed LC frequencies and subset composition in the oral mucosa. Overall, LC frequencies and absolute numbers were comparable between CD11c-Ecad^DEL^ and control mice (Fig. 1d), indicating that E-cadherin is not required to maintain the total LC count in the tissue. Intriguingly, analysis of the subsets revealed a marked shift in LC distribution (Fig. 1e). We detected significantly fewer LC1 and more LC2 in the gingiva, buccal mucosa and tongue, whereas the number of moLC was less affected (Fig. 1f-h, Suppl.Fig 1b,d). Phenotypic analysis further demonstrated subset-specific effects on activation status. While MHCII expression was slightly increased on LC1, LC2 exhibited a significant reduction in surface MHCII levels (Fig. 1i,j). Taken together, these results demonstrate that E-cadherin expression is required to maintain the structural integrity and subset composition of LC in the oral mucosa.

**Figure 1:**
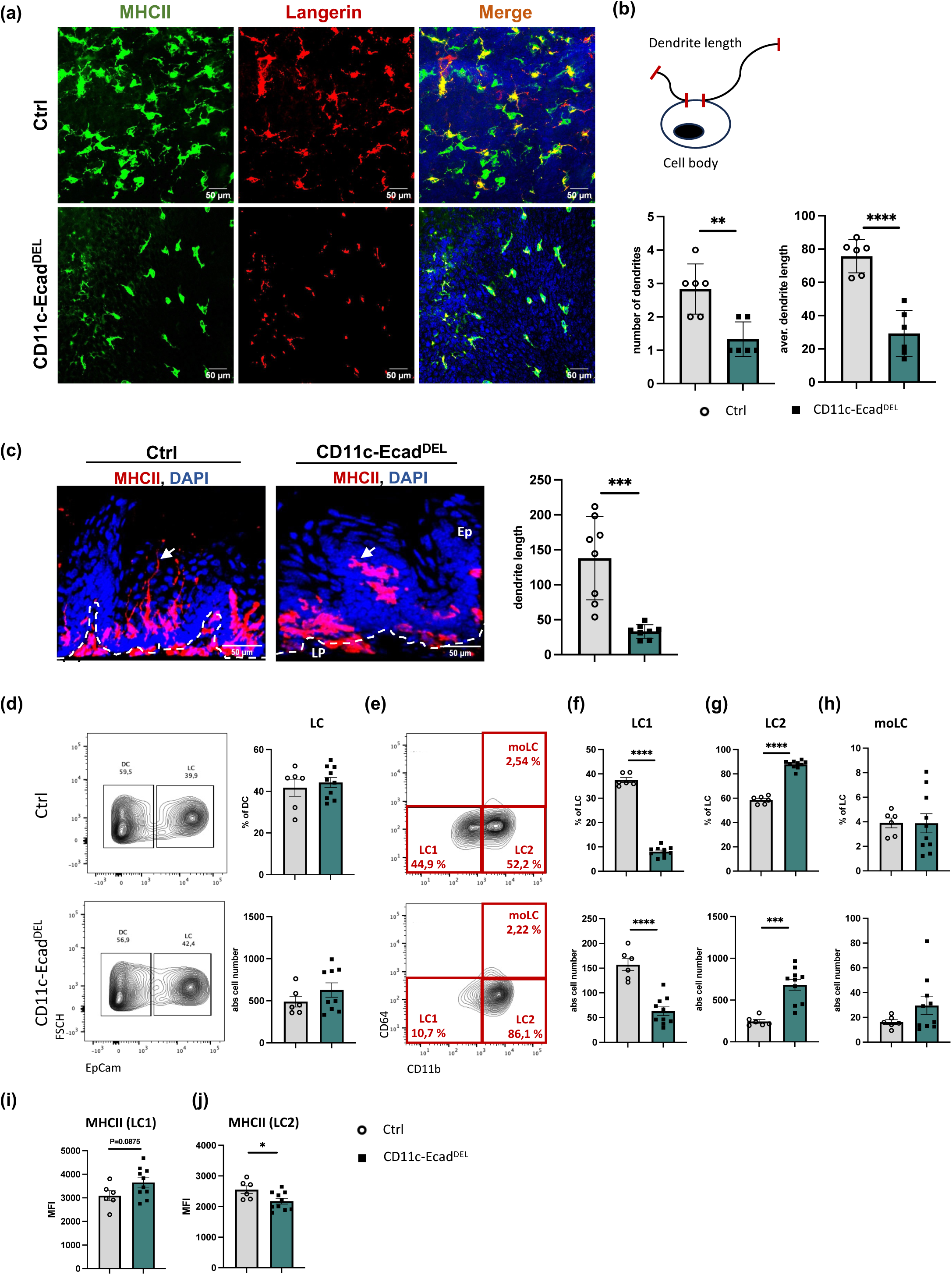
E-cadherin deletion in CD11c^+^ cells alters LC morphology and homeostasis in the oral mucosa. (a) Representative immunofluorescence images of buccal mucosa epithelial sheets from control and CD11c-Ecad^DEL^ mice, stained with antibodies against MHCII (green), Langerin (red), and DAPI (blue). (b) Quantification of LC morphology, including number of dendrites and dendrite length. (c) Representative tongue cross-sections from control and CD11c-Ecad^DEL^ mice, stained with antibodies against MHCII (red) to visualize LC dendrites and with DAPI (blue). White dotted lines represent the basal membrane. The graph shows the quantification of dendrite length. (d-j) Flow cytometric analysis of buccal mucosal LC subsets. (d) Representative flow cytometry plots identifying EPCAM^+^ LC, and (e) LC subsets pre-gated on CD45^+^CD11c^+^MHCII^+^ cells. Graphs show frequencies and absolute cell numbers of (d) LC, (f) LC1, (g) LC2, and (h) moLC. (i,j) Mean fluorescent intensity of MHCII on buccal LC1 and LC2 subsets. Data are presented as the mean ± SEM and are representative of two to three independent experiments. Statistical significance was determined using the Student’s t-test (* P < 0.05; ** P < 0.01; *** P < 0.005).

### E-cadherin is dispensable for LC migration under steady-state and inflammatory conditions

To determine whether the changes in LC subset composition observed in CD11c-Ecad^DEL^ mice were attributable to altered migration, we analyzed LC trafficking to the draining LN. First, we assessed the migratory potential of oral LC by determining CCR7 expression on LC. CCR7 levels were comparable between CD11c-Ecad^DEL^ and control mice – regarding both on the total LC population and on individual LC subsets – in both the tissue and the draining LN (Fig. 2b,c). In addition, flow cytometric analysis revealed comparable frequencies and absolute cell numbers of the total LC population as well as LC1 and LC2 in the draining LN of CD11c-Ecad^DEL^ and control mice (Fig. 2a,d), indicating that steady-state oral LC migration is not affected by the loss of E-cadherin.

**Figure 2:**
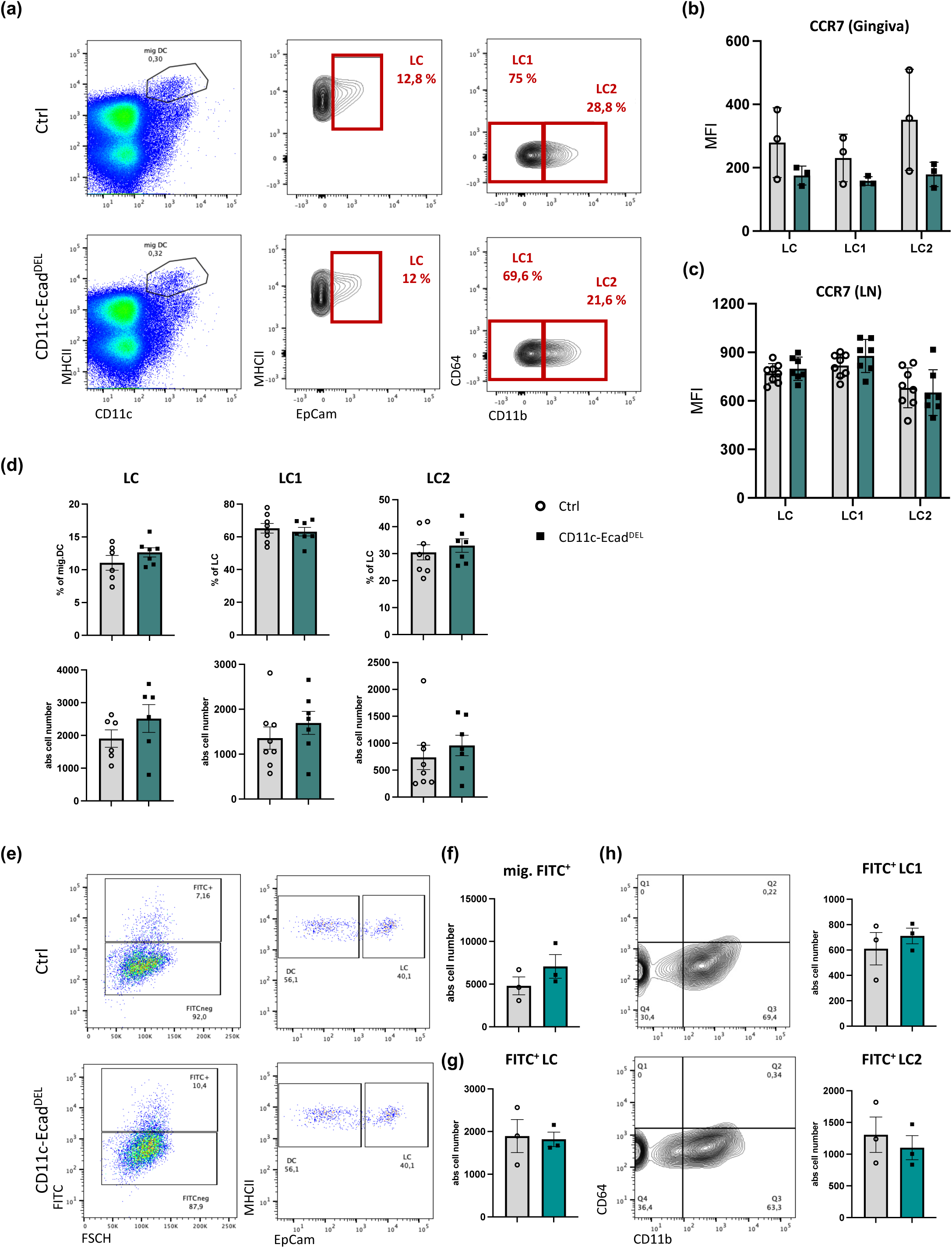
Loss of E-cadherin does not affect LC migration at steady-state and under inflammatory conditions. (a) Flow cytometric analysis of steady-state oral mucosa-draining LN from CD11c-Ecad^DEL^ and control mice. CCR7 expression on LC and LC subsets in the Gingiva (b) and LN (c), represented by mean fluorescent intensity. (d) Graphs show frequencies and absolute cell numbers of all LC, LC1, and LC2. (e) The gingiva was painted with FITC-solution, and three days later, draining cervical LN were collected and analyzed by flow cytometry. Representative FACS plots gated on CD11c^+^MHCII^hi^ tissue-derived dendritic cells. Furthermore, FITC-labeled EpCam^+^ LC, representing migratory LC, were detected in the LN. Absolute cell number of (f) FITC⁺ migratory cells and (g) FITC^+^ LC. (h) Representative FACS plots and absolute cell numbers of LC subsets in the draining LN of CD11c-Ecad^DEL^ and control mice. Data are presented as the mean ± SEM and are representative of two to three independent experiments. Statistical significance was determined using Student’s t-test (* P < 0.05; ** P < 0.01; *** P < 0.005).

Since inflammatory signals can enhance LC emigration from the tissue, we next assessed migration under inflammatory conditions using a FITC-painting model. By applying FITC to the gingiva, we were able to track migrating cells originating from the tissue to the draining cervical LN. Three days after FITC exposure, FITC⁺ migratory LC were detected within the CD11c⁺MHCII^hi^ compartment (Fig 2e,f). Notably, both the frequency and absolute number of FITC-labeled LC were comparable between CD11c-Ecad^DEL^ and control mice (Fig. 2g). Subset analysis further confirmed that migration of LC1 and LC2 populations remained unaffected (Fig. h). Collectively, these findings demonstrate that E-cadherin–mediated adhesion is not required to control oral LC migration to draining LN, neither under steady-state conditions nor during inflammation.

### E-cadherin deficiency in LC induces oral dysbiosis

The absence of E-cadherin resulted in impaired dendrite formation and extension towards the epithelial surface (Fig. 1), which may compromise the ability of LC to efficiently interact with the resident microbiota (dental biofilm). This, in turn, may lead to dysbiosis and inflammation (Capucha, Koren et al. 2018). Therefore, we analyzed the oral microbiota of CD11c-Ecad^DEL^ and control mice. Quantification of the total bacterial load by 16S qPCR revealed a trend towards increased bacterial abundance in the oral cavity of CD11c-Ecad^DEL^ mice compared to controls (Fig. 3a). At the genus level, we observed a significant reduction in *Lactobacillus* spp., whereas *Streptococcus* spp. showed an increased abundance (Fig. 3b, c). To resolve these changes at the community level, we performed 16S rRNA gene amplicon sequencing of oral and fecal samples. The oral microbiota of control mice was dominated by *Muribacter muris*, together with *Streptococcus spp*. In contrast, CD11c-Ecad^DEL^ mice displayed a more evenly distributed community characterized by reduced *Muribacter*, expanded *Streptococcus*, and the emergence of multiple lower-abundance taxa (Fig. 3d). Consistent with this, both the observed richness and Shannon diversity in the oral microbiota of CD11c-Ecad^DEL^ mice were significantly increased (Fig. 3e). On the other hand, the fecal microbiota of the same animals was comparable between genotypes in composition (Fig. 3f) and unchanged in alpha diversity (Fig. 3g). Beta-diversity analysis based on generalized UniFrac distances revealed a clear separation of oral communities by genotype along the first principal coordinate (59.4 % of variance) (Fig. 3h). Thus, loss of E-cadherin on LC drives a pronounced, orally restricted remodeling of the microbiota.

**Figure 3:**
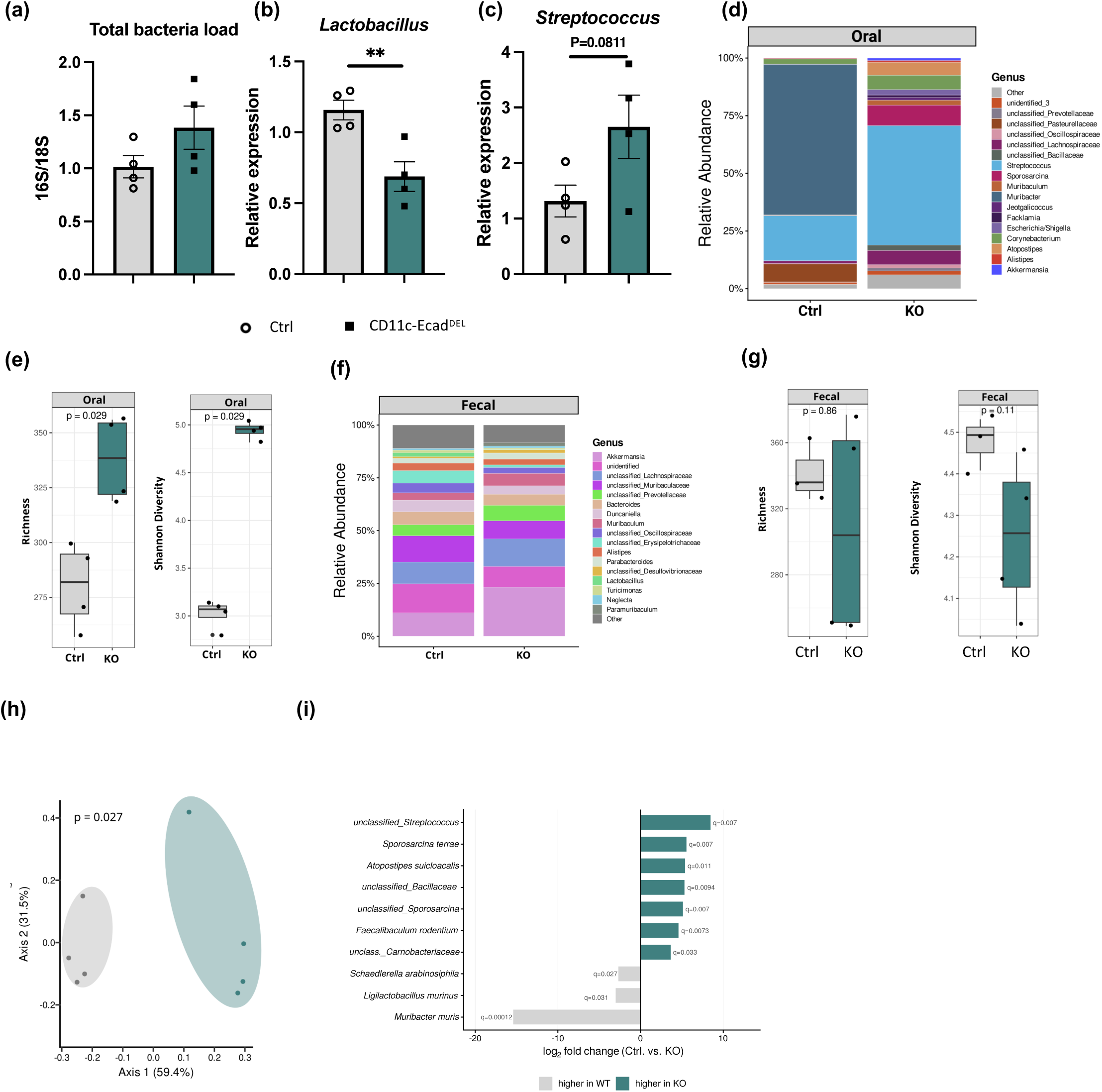
E-cadherin deficiency on LC induces oral dysbiosis. (a) Quantification of total bacterial load in oral samples from CD11c-Ecad^DEL^ and control mice by qPCR analysis of bacterial 16S rRNA gene copies. (b) Relative expression of *Lactobacillus spp.* and (c) *Streptococcus spp.* in oral swabs of CD11c-Ecad^DEL^ and control mice. (d) Relative abundance of the oral microbiota at the genus-level, determined by 16S rRNA gene amplicon sequencing. (e) Oral alpha diversity (observed richness and Shannon diversity). Data are presented as boxplots (median, interquartile range, and individual mice), n = [4] per group. (f) Relative abundance of the fecal microbiota at the genus-level. (g) Fecal alpha diversity (observed richness and Shannon diversity). (h) Beta diversity of the oral microbiota, presented as principal coordinates analysis (PCoA) of generalized UniFrac distances; axis labels indicate the proportion of variance explained. (i) Differentially abundant species in the oral microbiota, identified by LinDA, displayed as log2-fold change (control vs. CD11c-Ecad^DEL^); colored bars denote genera significant at padj < 0.05, with q-values indicated. Statistical significance: (a-c) unpaired two-tailed t-test and data are presented as mean ± SEM ; (e, g) Wilcoxon rank-sum test; (h) PERMANOVA; (i) LinDA with Benjamini–Hochberg correction. **p < 0.01; exact p-values are given where applicable. PCR data are presented as mean ± SEM.

Differential abundance analysis (LinDA) identified the genera underlying this divergence (Fig. 3i). The oral microbiota of CD11c-Ecad^DEL^ mice was reduced of *Muribacter muris* (q = 1.2 × 10⁻⁴), *Ligilactobacillus murinus* (q = 0.031), and *Schaedlerella arabinosiphila*, , while *Streptococcus spp.*, *Atopostipes suicloacalis*, *Sporosarcina spp.*, unclassified Bacillaceae, *Faecalibaculum rodentium*, and unclassified Carnobacteriaceae were enriched (padj < 0.05). Hence, E-cadherin deficiency was characterized by a marked reduction in the host-adapted commensal core that defines the healthy community – most prominently the dominant *Muribacter* – together with a compensatory expansion of *Streptococcus* species and multiple, largely low-abundance opportunistic taxa. The reduction of *Ligilactobacillus murinus* corroborated the qPCR-based loss of Lactobacillus spp. and is consistent with a depletion of protective lactic-acid bacteria (Tanwar, Gnanasekaran et al. 2026).

To assess whether this dysbiosis persisted with age, we analyzed the oral and fecal microbiota of 32-week-old mice (Suppl. Fig. 2). The oral dysbiosis was still detectable at this later timepoint but clearly less pronounced than in young mice: oral Shannon diversity remained significantly increased in CD11c-Ecad^DEL^ mice, with richness showing a corresponding trend (Suppl. Fig. 2b), and oral communities remained significantly separated by genotype (Suppl. Fig. 2e). However, the number of differentially abundant genera was markedly reduced relative to young mice, with only the decrease of *Muribacter muris* and unclassified Pasteurellaceae as well as the expansion of *Streptococcus spp.* remaining significant (Suppl. Fig. 2f). The fecal microbiota in these mice was again unaffected in both composition and diversity (Suppl. Fig. 2c, d). Overall, the genotype-dependent differences in the oral microbiota became less pronounced with age: the depletion of the *Muribacter* commensal core represented a stable, age-independent signature of the dysbiosis, whereas the increased diversity and the broader expansion of opportunistic taxa seen at 8 weeks were attenuated at 32 weeks.

Taken together, these data demonstrate that E-cadherin expression on LC is required to maintain oral microbial homeostasis: its loss results in a compositionally distinct, more diverse oral community marked by collapse of the commensal *Muribacter*/*Lactobacillus* core and expansion of opportunistic taxa – a dysbiotic state that is restricted to the oral mucosa and persists, in its core features, into aged animals.

### E-cadherin deficiency in LC induces inflammatory reprogramming in the epithelium and spontaneous alveolar bone loss

Dysbiosis can lead to persistent inflammation, which in turn drives alveolar bone loss, a hallmark of periodontal disease. To assess the impact of E-cadherin deficiency in LC on immune and tissue homeostasis under steady-state conditions, we performed bulk RNA sequencing of the gingival epithelium of 30-week-old CD11c-Ecad^DEL^ and control mice (Fig. 4a-c). Unsupervised clustering revealed a clear separation between CD11c-Ecad^DEL^ and control samples (Fig. 4a), indicating substantial transcriptional reprogramming in the absence of E-cadherin. Differential gene expression analysis demonstrated a predominance of upregulated genes in CD11c-Ecad^DEL^ mice, as illustrated by volcano plot analysis (Fig. 4b). Gene set enrichment analysis (GSEA) revealed a significant enrichment of immune-related and inflammation-associated pathways, including NF-κB, TNF, IL-17, Toll-like receptor, and MAPK signaling (Fig. 4c). Consistent with the transcriptomic profile, Luminex multiplex analysis of the gingiva confirmed increased concentrations of pro-inflammatory cytokines in CD11c-Ecad^DEL^ mice compared to control animals. In particular, IL-17A, a pro-inflammatory cytokine implicated as a critical driver of alveolar bone loss in periodontitis (Dutzan, Kajikawa et al. 2018, Hajishengallis and Lamont 2021), was significantly elevated (Fig. 4d). These data indicate a shift towards a Th17-dominated inflammatory environment. In line with this, flow cytometric analysis of the T cell compartment in the gingiva revealed an expansion of both αβ and γδ T cells (Fig. 4e,f), indicating enhanced T-cell–mediated immune activation in the absence of E-cadherin on LC. Intriguingly, this inflammatory phenotype in the gingival tissue was age-dependent, as young mice (8–12 week-old) displayed comparable cytokine levels and T-cell frequencies between CD11c-Ecad^DEL^ and control mice (SupplFig. 3a-d). Given the established association between dysbiosis, chronic gingival inflammation, and alveolar bone loss, we subsequently assessed bone loss in 50-week-old CD11c-Ecad^DEL^ and control mice. Expression of *RankL*, a key factor in alveolar bone resorption, was significantly increased in aged CD11c-Ecad^DEL^ mice (Fig. 4g), while only a slight trend was observed in young animals (12 week-old, SupplFig. 3e). Consistent with this finding, micro-computed tomography (µCT) analysis revealed a significant increase in alveolar bone loss in CD11c-Ecad^DEL^ mice under steady-state conditions compared to controls (Fig. 4h). Collectively, these results demonstrate that the lack of E-cadherin on LC promotes a pro-inflammatory tissue environment characterized by enhanced T cell infiltration, elevated cytokine production, and spontaneous bone loss.

**Figure 4:**
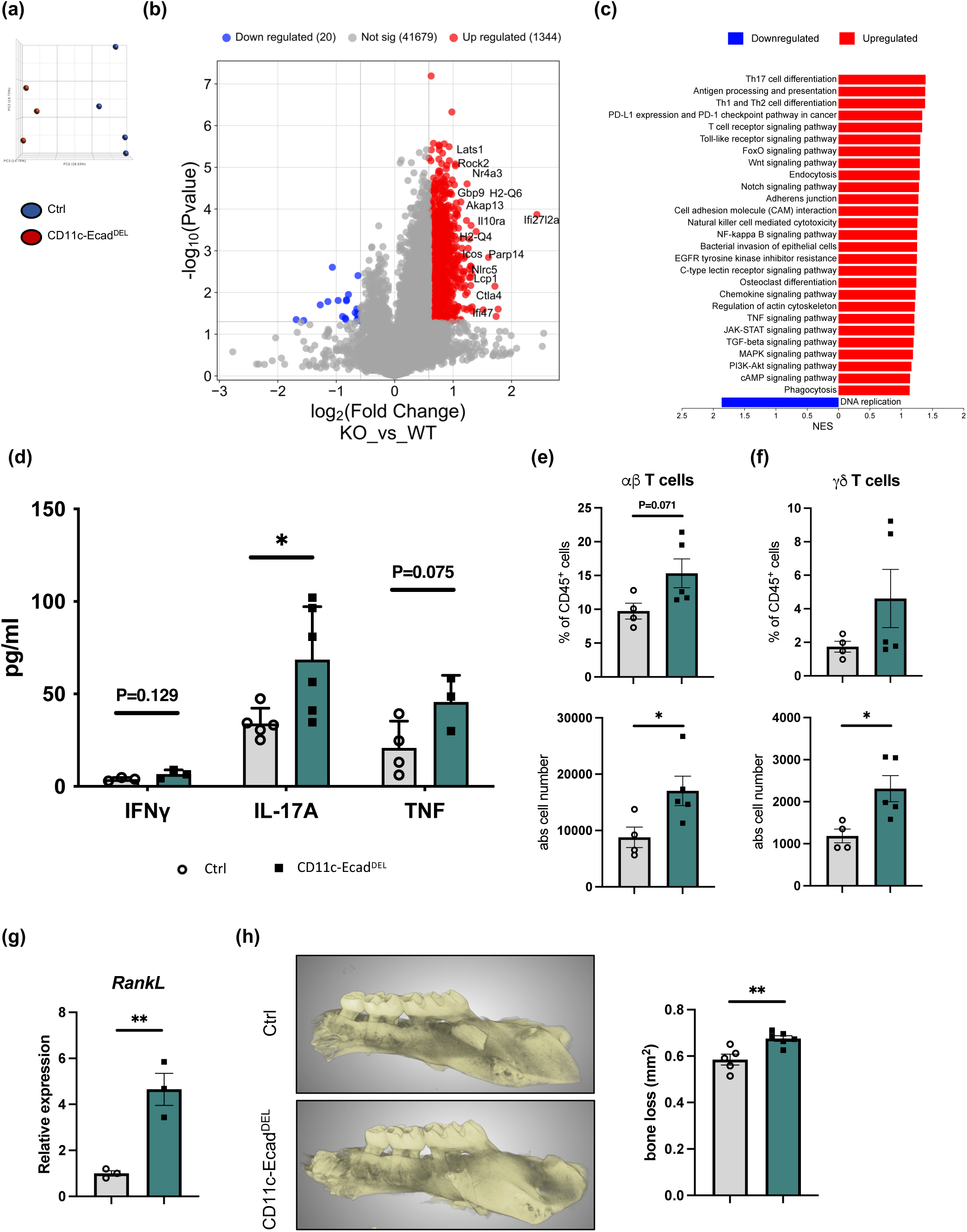
E-cadherin deficiency in LC drives inflammatory reprogramming and spontaneous alveolar bone loss. (a-c) Bulk RNA transcriptome profile of gingiva epithelium of CD11c-Ecad^DEL^ versus control mice. (a) Principal component analysis (PCA) showing distinct clustering of CD11c-Ecad^DEL^ compared to control samples. (b) Volcano plot of differentially expressed genes highlighting significantly upregulated and downregulated genes in CD11c-Ecad^DEL^ versus Ctrl mice. Cutoff value: fold change 1.5 and -1.5, respectively; p-value < 0.05. (c) KEGG pathway enrichment analysis of differentially expressed genes with normalized enrichment scores (NES) for pathways upregulated (red) and downregulated (blue) in CD11c-Ecad^DEL^ versus control samples. (d) Gingiva IFNɣ, IL-17A and TNF cytokine levels as determined by Luminex multiplex assay. (e-f) Flow cytometric analysis and quantification of gingival αβ and ɣδ T cells of 25 week-old control and CD11c-Ecad^DEL^ mice. (g) Relative expression of *RankL* in the gingiva of control and CD11c-Ecad^DEL^ mice, measured by qRT-PCR. (h) Representative μCT images of the jaw bones from control and CD11c-Ecad^DEL^ mice. Distance between the cementoenamel junction (CEJ) and alveolar bone crest (ABC) was used to determine bone loss. The graph shows the quantification of bone loss. Data represent the mean values ± SEM, where each dot denotes an individual mouse. Data are representative of two to three independent experiments. Statistical significance was determined using Student’s t-test (* P < 0.05; ** P < 0.01).

### Lack of E-cadherin on LC leads to increased alveolar bone loss in ligature-induced periodontitis

To determine the functional consequences of altered LC homeostasis and dendrite formation during inflammation, we assessed disease severity in a ligature-induced periodontitis (LIP) model. To this end, ligatures were applied around the first molars of CD11c-Ecad^DEL^ and control mice for up to 7 days to induce local oral inflammation and gingival immune responses were evaluated by flow cytometry (Nogueira, Nokhbehsaim et al. 2022, Rath-Deschner, Nogueira et al. 2022, Netanely, Barel et al. 2024). This analysis revealed a marked increase in inflammatory cell infiltrates in CD11c-Ecad^DEL^ mice, characterized by elevated frequencies of Ly6G^hi^ neutrophils and Ly6C^hi^ monocytes in the gingival tissue compared to controls (Fig. 5a-c). This enhanced cellular infiltration was accompanied by increased production of pro-inflammatory cytokines (Fig. 5d), MCP1 and RANKL (Fig. 5e). Consistent with the increased inflammatory environment, µCT analysis demonstrated higher alveolar bone loss in CD11c-Ecad^DEL^ mice compared to control animals (Fig. 5f,g). This indicates a protective role of E-cadherin–dependent LC interactions in periodontal tissue integrity. Deletion of E-cadherin in LC exacerbates periodontal inflammation and bone loss during LIP, thus highlighting a critical role for LC-mediated immune regulation in limiting tissue-destructive responses during periodontitis.

**Figure 5:**
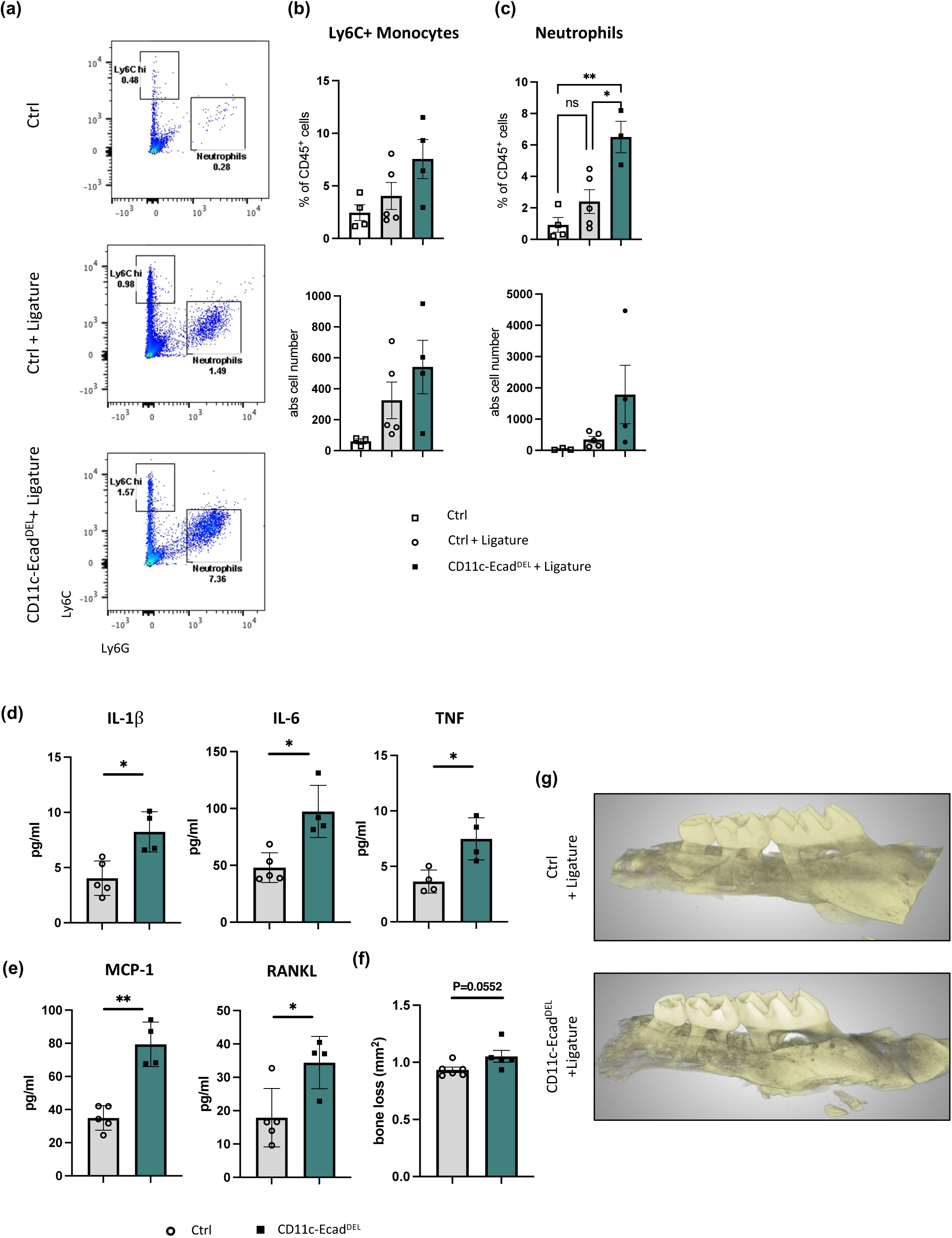
Absent E-cadherin on LC exacerbates inflammation and alveolar bone loss in ligature-induced periodontitis. (a) Representative flow cytometry plots of gingival immune cells from control and CD11c-Ecad^DEL^ mice depicting Ly6C^hi^ monocytes and Ly6G^hi^ neutrophils. Quantification of (b) Ly6C^hi^ monocytes and (c) Ly6G^hi^ neutrophils in the gingiva presented as frequencies and absolute cell numbers. (d) Gingiva IL-1β, IL-6 and TNF cytokine levels and (e) MCP-1 and RANKL as determined by Luminex multiplex assay. (f) Quantification of alveolar bone loss in CD11c-Ecad^DEL^ compared to control mice. (g) Representative three-dimensional µCT reconstructions of maxillary alveolar bone from control and CD11c-Ecad^DEL^ mice. The distance between the cementoenamel junction (CEJ) and alveolar bone crest (ABC) was used to determine bone loss. Data are presented as mean ± SEM, with each dot representing an individual mouse. Data are representative of two independent experiments. Statistical significance was determined using Student’s t-test (* P < 0.05; ** P < 0.01; *** P < 0.001).

## Discussion

In this study, we identify E-cadherin as a critical regulator of oral LC morphology and mucosal immune homeostasis. Using CD11c-specific E-cadherin–deficient mice, we demonstrate that loss of E-cadherin profoundly alters LC morphology and subset composition, disrupts oral microbial homeostasis, promotes inflammatory reprogramming of the gingival tissue, and exacerbates alveolar bone loss under both steady-state and inflammatory conditions. Together, our findings establish a previously unrecognized role for E-cadherin in shaping the interplay between LC, oral epithelium and microbiota, and thereby controlling periodontal homeostasis.

One significant finding of our study is the profound alteration of LC morphology in the absence of E-cadherin. Oral LC lacking E-cadherin exhibit a rounded phenotype with markedly reduced dendrite formation and impaired extension towards the epithelial surface. LC dendrites are essential for immune surveillance and antigen sampling at barrier tissues (Kubo, Nagao et al. 2009, Kaplan 2017). In the skin, LC dynamically extend dendrites through epithelial tight junctions to capture antigens while maintaining epithelial integrity (Kubo, Nagao et al. 2009). Our findings suggest that similar mechanisms operate in the oral mucosa and critically depend on E-cadherin–mediated epithelial interactions. These observations are consistent with our previous work showing that E-cadherin–deficient epidermal LC also exhibit reduced dendritic processes despite preserved tissue residency (Brand, Diener et al. 2020). While this does not obviously affect skin homeostasis, the consequences of failing dendrite formation in the oral mucosa are substantially more pronounced, likely reflecting the constant microbial exposure of the more delicate oral barrier.

E-cadherin is a calcium-dependent adhesion molecule and central component of adherens junctions in epithelial tissues (van Roy and Berx 2008). Since the seminal observation that LC attach to keratinocytes via homophilic E-cadherin binding (Tang, Amagai et al. 1993), E-cadherin has been widely considered essential for LC retention within stratified epithelia (Jakob and Udey 1998). Notably, here we show that E-cadherin deficiency does not affect total LC numbers in the oral mucosa and draining LN, indicating that E-cadherin is dispensable for LC maintenance and tissue retention under steady-state conditions. Furthermore, LC migration to LN is not changed under steady-state or inflammatory conditions, which is supported by similar CCR7 expression on LC in the oral mucosa and draining LN. This is in line with findings in epidermal LC, which demonstrate a preserved LC network and turnover after conditional deletion of E-cadherin (Brand, Diener et al. 2020). Our data further support the notion that LC residency can be maintained independently of E-cadherin, potentially through compensatory adhesion pathways or tissue-derived survival signals.

Given the well-established bidirectional relationship between mucosal LC and the local microbiota, where microbial signals shape oral LC development and LC in turn contribute to tissue homeostasis and protection from immunopathology (Capucha, Koren et al. 2018), disrupted LC morphology may have important functional consequences for oral immune regulation. Despite unchanged total LC frequencies, we observe a shift in LC subset composition characterized by reduced LC1 and increased LC2 populations. In addition, phenotypic analyses reveal subset-specific changes in activation status. While LC1 display slightly increased MHCII expression, LC2 exhibit significantly reduced MHCII surface levels, indicating that loss of E-cadherin adhesion signaling may affect LC subsets in distinct ways. Whether these alterations reflect differences in epithelial interactions or responsiveness to local environmental cues remains to be determined.

Previous studies identified oral LC subsets with distinct developmental origins and potentially specialized functions (Capucha, Mizraji et al. 2015). Notably, germ-free mice display reduced CD103⁺ LC1 frequencies, suggesting that this subset is particularly dependent on microbial-derived signals (Capucha, Koren et al. 2018). Our findings therefore raise the possibility that impaired dendrite formation compromises the ability of, in particular, LC1 to efficiently sense microbial cues at the epithelial surface, thereby affecting their local maintenance or differentiation. In contrast, LC2 appear to be less dependent on E-cadherin, indicating differential niche requirements between LC subsets. Whether these subsets indeed exert distinct regulatory or pro-inflammatory functions during periodontal disease remains an important question for future studies.

Another key finding of this study is that defective LC morphology is associated with substantial oral microbial dysbiosis. CD11c-Ecad^DEL^ mice exhibit altered microbial composition and elevated microbial diversity. Notably, these changes were not mirrored in the fecal microbiota, which remained largely unaffected, suggesting that loss of E-cadherin on LC selectively impacts the oral microbial niche rather than inducing a broad systemic microbiota alteration. Previous work demonstrated that prolonged LC depletion results in increased oral bacterial load and aggravated periodontal pathology (Capucha, Koren et al. 2018). Our data now extend these observations by showing that even in the presence of LC, disruption of LC dendrite organization, and hence impaired interaction with the epithelium and oral microbiota, is sufficient to induce dysbiosis. At the community level, the oral microbiota of control mice is dominated by a single commensal species, *Muribacter muris*, which is strongly reduced in CD11c-Ecad^DEL^ mice alongside a decrease in *Ligilactobacillus murinus (a species reclassified from the former Lactobacillus genus in 2020)* (Zheng, Wittouck et al. 2020), with a compensatory expansion of multiple low-abundance taxa. *Muribacter muris* is a host-adapted murine commensal that mediates colonization resistance against non-typable *Haemophilus influenzae* (Nicklas, Bisgaard et al. 2015, Granland, Scott et al. 2020); its loss therefore reflects the breakdown of a dominant protective commensal rather than a neutral compositional change. The accompanying rise in richness and evenness should not be read as beneficial but as consistent with collapse of the commensal-dominated community and loss of colonization resistance. Mechanistically, impaired epithelial surface access may reduce microbial sampling and alter LC-mediated regulation of immune tolerance towards commensals, relaxing the host control that normally constrains oral community composition. The observed reduction in *Lactobacillus spp*. – recovered both by targeted qPCR and, at the sequence level, as a depletion of the salivarius-clade species *Ligilactobacillus murinus* – together with expansion of *Streptococcus spp.* further supports a shift towards a dysbiotic microbial community associated with inflammatory conditions and periodontal disease (Dutzan, Kajikawa et al. 2018, Hajishengallis and Lamont 2021). *Lactobacillus* species are generally considered beneficial commensals that contribute to oral homeostasis through antimicrobial activity, competitive exclusion of pathogenic bacteria, and immunomodulatory functions, including suppression of pro-inflammatory cytokines such as TNF, IL-1β, and IL-17 (Koll-Klais, Mandar et al. 2005, Szkaradkiewicz, Stopa et al. 2014, Kobayashi, Kobayashi et al. 2017). In experimental periodontitis, oral administration of Lactobacillus strains reduces periodontal inflammation and alveolar bone loss, supporting their protective role in maintaining mucosal immune balance (Teughels, Durukan et al. 2013, Kobayashi, Kobayashi et al. 2017, Invernici, Salvador et al. 2018). In contrast, expansion of oral *Streptococcus* species has frequently been linked to inflammatory remodeling of the oral microbiota and enhanced immune activation (Abranches, Zeng et al. 2018, Lamont, Koo et al. 2018). Although certain commensal Streptococci contribute to colonization resistance under physiological conditions, excessive accumulation of *Streptococcus spp.* can promote epithelial activation and persistent inflammatory signaling (Kreth, Merritt et al. 2009, Lamont, Koo et al. 2018). In particular, oral Streptococci have been shown to stimulate production of IL-1β, IL-6, TNF, and IL-17-associated cytokines, thereby contributing to chronic mucosal inflammation and periodontal tissue destruction (Dutzan, Kajikawa et al. 2018, Hajishengallis and Lamont 2021). Increased abundance of inflammatory streptococcal communities has also been reported in human gingivitis and periodontitis and correlates with dysbiotic microbial shifts and disease severity (Griffen, Beall et al. 2012, Abusleme, Dupuy et al. 2013). Several of the expanded genera carry opportunistic, pro-inflammatory potential, notably Carnobacteriaceae, which comprise the nutritionally variant streptococci *Granulicatella* and *Abiotrophia*, recognized agents of orally seeded endocarditis (Tellez, Ambrosioni et al. 2018); their outgrowth offers a plausible substrate for the IL-17-associated inflammation. This expansion is nonetheless heterogeneous and not uniformly pro-inflammatory, also including environmental spore-formers (*Sporosarcina*, unclassified Bacillaceae) and the anti-inflammatory gut commensal *Faecalibaculum* rodentium (Abe and Hajishengallis 2013, Zagato, Pozzi et al. 2020), whose enrichment more likely reflects oral–fecal exchange. At genus-level 16S resolution these data therefore argue against a single driver and instead for a loss of host-imposed selection that destabilizes the community and permits opportunistic, partly inflammatory, taxa to expand.

Intriguingly, the core of this dysbiosis persists into aged animals. Although the increased diversity and the expansion of low-abundance opportunists are attenuated by 32 weeks, the two dominant shifts that define the community remain: the depletion of the *Muribacter* commensal core and the reciprocal expansion of *Streptococcus spp.* are maintained at both timepoints. The dysbiosis therefore does not resolve with age but stabilizes into a persistent state characterized by the loss of the dominant protective commensal and a sustained streptococcal predominance, implying that the sustained loss of *Muribacter*-mediated colonization resistance together with persistent streptococcal overgrowth drives disease progression. Consistent with this dysbiotic phenotype, transcriptomic and immunological analyses reveal substantial inflammatory reprogramming in the gingival tissue of CD11c-Ecad^DEL^ mice. RNA sequencing identifies an enrichment of NF-κB, TNF, IL-17, Toll-like receptor, and MAPK signaling pathways, all of which are strongly associated with periodontal inflammation and bone destruction (Hajishengallis 2014, Moutsopoulos and Konkel 2018). In particular, the pronounced increase in IL-17A production and expansion of αβ and γδ T cells indicate a Th17-skewed inflammatory environment. IL-17 signaling plays a central role in periodontal pathology by promoting neutrophil recruitment, inflammatory cytokine production, and osteoclastogenesis including bone loss through induction of RANKL expression (Dutzan, Kajikawa et al. 2018, Hajishengallis and Lamont 2021). The expansion of γδ T cells is particularly intriguing, as these cells are potent early producers of IL-17 at barrier tissues and have been implicated in mucosal inflammatory diseases (Akitsu and Iwakura 2018).

Notably, the inflammatory alterations in CD11c-Ecad^DEL^ mice are age-dependent and accompanied by spontaneous alveolar bone loss in older animals. This suggests that persistent disruption of LC–microbiota interactions gradually drives inflammatory pathology over time. Age-associated periodontal inflammation has previously been linked to persistent low-grade immune activation, dysbiotic microbial shifts, and progressive osteoclastogenic responses within gingival tissues (Ebersole, Steffen et al. 2008, Hajishengallis 2010). Increased *RankL* expression in aged mice further supports enhanced osteoclastogenic activity as a mechanism underlying bone destruction (Kawai, Matsuyama et al. 2006, Bostanci, Abe et al. 2019). Persistent low-grade dysbiosis may therefore progressively amplify inflammatory responses and tissue damage over time (Lamont, Koo et al. 2018, Hajishengallis, Chavakis et al. 2020). Current concepts propose that periodontitis is not caused solely by direct pathogen-mediated tissue injury but rather results from dysregulated host immune responses to a dysbiotic microbiota (Hajishengallis 2014, Hajishengallis and Lamont 2014). Our findings are consistent with this model and suggest that defective LC-mediated epithelial surveillance in the absence of E-cadherin / dendrites initiates chronic dysbiosis that progressively drives inflammatory pathways and alveolar bone destruction during aging.

Finally, E-cadherin deficiency markedly exacerbates disease severity in ligature-induced periodontitis. CD11c-Ecad^DEL^ mice display increased recruitment of neutrophils and inflammatory monocytes together with enhanced cytokine production and alveolar bone destruction. Experimental ligature-induced periodontitis reproduces key pathological features of human periodontal disease, including microbial dysbiosis, inflammatory leukocyte infiltration, osteoclast activation, and progressive bone loss (Abe and Hajishengallis 2013, de Molon, Park et al. 2018, Marchesan, Girnary et al. 2018). In this model, excessive neutrophil accumulation and inflammatory monocyte recruitment are central drivers of periodontal tissue destruction through release of reactive oxygen species, proteolytic enzymes, and osteoclastogenic mediators (Hajishengallis 2015, Fine, Chadwick et al. 2021). Consistent with this, increased infiltration of neutrophils and monocytes in CD11c-Ecad^DEL^ mice is associated with elevated inflammatory cytokine production, including IL-17- and TNF-associated pathways known to promote periodontal pathology and osteoclastogenesis (Dutzan, Kajikawa et al. 2018, Hajishengallis and Lamont 2021). Our findings therefore indicate that intact LC-epithelial interactions are critical for limiting excessive inflammatory responses during periodontal challenge. Given the preserved migratory capacity of E-cadherin–deficient LC, the enhanced pathology likely reflects defective local immune regulation within the gingival epithelium rather than altered antigen transport to LN and T-cell priming. Impaired dendrite formation and reduced epithelial surface sampling by E-cadherin–deficient LC may therefore compromise local regulation of commensal-host interactions, resulting in exaggerated inflammation during periodontal challenge. This interpretation is consistent with emerging evidence that tissue-resident antigen-presenting cells function not only as migratory initiators of adaptive immunity but also as critical local regulators of barrier tissue homeostasis and inflammation (Naik, Bouladoux et al. 2012, Clausen and Stoitzner 2015, Moutsopoulos and Konkel 2018). Our data establish a strong association between altered LC morphology, dysbiosis, and inflammation.

A significant conceptual advancement of our study is that it allows the analysis of the role of morphological dynamics for the function of LC independent of their tissue residency or migration. Most previous studies investigating oral LC used depletion models, in which LC are absent from the tissue (Arizon, Nudel et al. 2012, Capucha, Koren et al. 2018). In contrast, CD11c-Ecad^DEL^ mice uniquely preserve LC numbers and migratory capacity despite profound defects in dendrite formation. E-cadherin–deficient LC remained present within the oral epithelium and migrated normally to draining LN under both steady-state and inflammatory conditions. Thus, the primary defect is not the absence of LC but the loss of dendrite-mediated epithelial surveillance. This is particularly relevant because it demonstrates that the mere presence of LC within the epithelium is insufficient to maintain oral homeostasis. Instead, proper dendrite formation and epithelial surface interaction appear to be critical for controlling the oral microbiota and limiting inflammatory pathology.

In conclusion, our study identifies E-cadherin as a critical regulator of oral LC morphology and mucosal immune homeostasis. E-cadherin is essential for proper dendrite formation and microbiota surveillance by oral LC, thereby limiting dysbiosis-driven inflammation and periodontal bone loss. These results open the perspective to identify LC-specific signaling pathways that could be targeted in the future to treat periodontitis.

## Material and Methods

### Mice

E-cadherin^fl/fl^ mice (Boussadia, Kutsch et al. 2002) were crossed to CD11c-Cre mice (Caton, Smith-Raska et al. 2007) to obtain conditional knockouts with specific deletion of E-cadherin in all CD11c^+^ cells (CD11c-Ecad^DEL^). All experiments were performed with 8–52 week-old, sex and age-matched animals compared to Cre-negative littermate controls. The mice were housed and treated in accordance with relevant laws and institutional guidelines of the Central Animal Facility of the University Medical Center Mainz.

### Cell preparation

Gingiva, buccal mucosa and tongue were mechanically disrupted and digested with 2mg/ml Collagenase II (Life technologies, Gibco) and 1mg/ml DNase I (Roche, Basel, Switzerland) in FACS buffer (PBS + 2% FCS) for 20min at 37°C with shaking at 1000rpm. 20µl EDTA (10mM) was added for 10min at 37°C. LN were mechanically disrupted and digested with 400U/ml Collagenase IV (Worthington Biochemical Corp., Lakewood, NJ) and 0.5U/ml DNase (Promega, Madison, WI) in RPMI for 30min at 37°C and 20µl EDTA (10mM) was added for 5min at RT. Subsequently, cells were filtered through 70µm cell strainers (BD Biosciences, San Jose, CA) to obtain single-cell suspensions. For experiments where epithelium and lamina propria were separated, tissues were pretreated with 2mg/ml Dispase II in FACS buffer for 20-40min at 37°C shaking. Epithelium was detached from the underlying lamina propria under a dissecting microscope, and both parts were processed as described above.

### Flow cytometry

LN and oral mucosal cells were pre-incubated in FACS-buffer containing Fc-Block (Biocompare, South San Francisco, CA) for 10min and then surface-stained with various combinations of fluorescence-conjugated antibodies at 4°C for 30min. For intracellular staining cells were fixed for 30min with 2% PFA, permeabilized with 0.1% Saponin and incubated with appropriate antibodies for 60min at 4°C. Flow cytometric acquisition was performed on a FACS Canto II (BD) and analyzed using FlowJo software (Treestar).

### Immunofluorescence microscopy

Oral tissues were prepared and stained as whole mount epithelial sheets or frozen cross-sections. Gingiva and tongue epithelial sheets were prepared by incubating/injecting tissue with 4mg/ml Dispase II (Roche) in PBS + 2% FCS and incubated for 10min at 37°C. Epithelium was carefully separated from the underlying lamina propria and fixed with 4% PFA for 10min. For frozen sections, whole tongue was excised, washed with PBS and fixed with 4% PFA for 2hrs and submerged in 30% sucrose solution overnight. Tissue was imbedded in tissue freezing medium (Leica Biosystems) and 20µm thick cross sections were prepared. Epithelial sheets and cross sections were stained as floating sections. Briefly, sections were washed with PBS, blocked in blocking buffer containing 15% FBS, 5% BSA and 0.3% Triton-X 100 in PBS for 3hrs at room temperature and incubated in fluorochrome-conjugated primary antibodies overnight. Sections were washed in PBS and mounted on slides with mounting medium containing 4,6-Diamidin-2-phenylindol (DAPI, Sigma). Images were acquired using a Leica TCS SP8 confocal microscope (Leica biosystems) and analyzed using Image J.

### Antibodies

The following anti-mouse monoclonal antibodies (clone) from BD Biosciences, BioLegend and eBioscience were used for flow cytometry: CD11c (N418), MHC-II (M5/114), CD45 (30-F11), CD11b (M1/70), CD103 (M290), CD64 (X54-5/7.1), CD24 (M1/69), Langerin (929F3), EpCam (G8.8), E-cadherin (24E10; Cell Signaling), CD40 (3/23), CD80 (16-10A1), CD86 (GL1), CCR7 (4B12), CD4 (L3T4), CD8α (53-6.7), TCRβ (H57-597), and TCRγδ (eBio GL3).

### Cytokine detection

Cell culture supernatant levels of IFNɣ, IL-17A and TNF were determined by Luminex Multiplex Assays according to the manufacturer’s instructions (ThermoFischer, Waltham, MA).

### Bacterial DNA analysis

To examine the absolute abundance of certain bacteria, DNA was prepared from oral swabs using Zymo Quick-DNA Fungal/Bacterial DNA Extraction Kit according to the manufacturer’s instructions, and then directly subjected to 16S RT-PCR analysis using designated primers (16sfw: AGA GTT TGA TCC TGG CTC; 16srev: TGC TGC CTC CCG TAG GAG T; 18sfw: CGG CTA CCA CAT CCA AGG AA; 18srev: GGG CCT CGA AAG AGT CCT GTA T; Lactobacillusfw: GGA AAC AG (A/G) TGC TAA TAC CG; Lactobacillusrev: ATC GTA TTA CCG CGG CTG CTG GCA; Streptococcusfw: CCT ACG GGA GGC AGC AGT AG; Streptococcusrev: CAA CAG AGC TTT ACG ATC CGA AA). The RT-qPCR reaction was performed using Power SYBR Green PCR Master Mix (Quanta-BioSciences IncTM). The samples were normalized to the 18S gene copies using the change in cycling threshold (DCT) method and calculated based on 2-DCT.

### Microbiota analysis

Oral microbiome samples were collected from mice using Copan minitip flocked swabs in 1ml eNAT® medium (Copan, Brescia, Italy). For fecal microbiota analysis, freshly collected stool pellets were resuspended in 1ml magiX PB1 microbiome stabilization buffer (microBIOMix, Regensburg, Germany). All samples were stored at –80°C and subsequently shipped on dry ice for microbiome sequencing. Samples were processed in a single batch. DNA was isolated of fecal and oral suspensions by bead-beating with Lysing Matrix Y (MP Biomedicals, USA) on a TissueLyser II (Qiagen, Germany) and purified on a MagNA Pure 96 system (Roche Diagnostics, Switzerland). Bacterial 16S rRNA gene copies were quantified by qPCR (LightCycler 480 II, primers 764F/907R, SYBR Green I Master; Roche Diagnostics) against full-length 16S rDNA standards (27F/1492R products cloned into pGEM-T Easy; Thermo Fisher Scientific, USA).The V4–V6 region was amplified with primers 341F and 1061R for oral and the V3–V4 hypervariable regions for fecal samples using primers 341F/815R. Bacterial 16S rRNA genes were amplified from 1 × 10^7^ 16S rRNA gene copies per sample. Barcoded amplicons were pooled, purified with MagSi-NGSPREP-PLUS beads (Steinbrenner Laborsysteme, Germany; 1:1.2 ratio), quantified with the Ion Library TaqMan™ Quantitation Kit, and sequenced on an Ion GeneStudio S5 Plus platform after Ion Chef templating with the Ion 520™ & Ion 530™ ExT Kit–Chef (Thermo Fisher Scientific). Raw data (Torrent Suite 5.18) were adapter/primer-trimmed and demultiplexed with cutadapt 4.4, quality-trimmed with Trimmomatic 0.39 (10-base sliding window, Q15, reads < 250 bp discarded), and filtered for > 5 expected errors with vsearch 2.28.1. Zero-radius operational taxonomic units (zOTUs) were defined (alpha 2, minimum five reads), chimeras removed with uchime3_denovo, and reads mapped at ≥ 98 % identity via usearch_global. Taxonomy was assigned in R 4.4.0 with IDTAXA (DECIPHER 2.26.0) against RDP version 19 trainingset (bootstrap 98 %, score threshold 40). A phylogenetic tree was built by alignment of zOTU sequences using FastTree 2.2.0. Alpha diversity was calculated as zOTU richness and Shannon diversity using mia 1.1.7. Beta diversity was assessed by generalized UniFrac analysis (GUniFrac package, weighting parameter alpha was set to 0.5) computed from a zOTU phylogenetic tree, followed by principal coordinates analysis; group differences were tested by PERMANOVA (pairwiseAdonis::pairwise.adonis). Differential abundance was analysed with MicrobiomeStat::linda (LinDA 1.1) at species level. All plots were generated with ggplot2 3.5.1. Raw sequencing data for this project have been deposited in the European Nucleotide Archive (ENA) under accession number PRJEB115995.

### RNA isolation and RT-PCR

For RNA isolation, the excised gingival tissues and tongues were homogenized in 500 µl TRI reagent (Sigma) using an electric homogenizer and RNA was extracted according to the manufacturer’s instructions. cDNA synthesis was performed using the QuantiTect Reverse Transcription Kit according to the manufacturer’s instructions (QIAGEN). RT-PCR reactions were performed using SYBR® Green PCR Master Mix and specific primers (*HPRT*fw: CGT CGT GAT TAG CGA TGA TG ; *HPRT*rev: TCC AAA TCC TCG GCA TAA TG ; *RankL*fw: TGT ACT TTC GAG CGC AGA GAT G ; *RankL*rev: AGG CTT GTT TGA TCC TCC TG). The samples were normalized to *HPRT* as control mRNA by change in cycling threshold (ΔCT) method and calculated based on 2-ΔΔCT.

### FITC painting

20 mg of FITC (Sigma) was dissolved in 100µl DMSO (Sigma), and the solution was diluted in acetone (1:1). Mice were anesthetized, and 40µl of the FITC-solution was carefully applied to the gingiva. After 48hrs draining cervical LN were collected, enzymatically digested to obtain a single-cell suspension and analyzed for the frequency of incoming FITC^+^ migratory dendritic cells by flow cytometry.

### Bulk RNA sequencing

RNA was purified using the RNeasy Plus Micro Kit (Qiagen) according to the manufacturer’s protocol. RNA was quantified with a Qubit Flex fluorometer (Thermo Fisher Scientific) and the quality was assessed on a Bioanalyzer 2100 (Agilent) using an RNA 6000 Pico chip (Agilent). Samples with an RNA integrity number (RIN) of ≥ 8 were sent to Novogene (Cambridge, UK) for library preparation (polyA approach) and sequencing. Sequencing was performed on Illumina’s NovaSeq 6000. Raw sequencing reads (approx. 30 million PE150 reads per sample) were preprocessed according to the Illumina standard protocol. Sequence reads were further processed using Qiagen’s CLC Genomics Workbench (v23.0.2) software with CLC’s default settings for RNAseq analysis. Reads were aligned to GRCm39 genome (assembly GRCm39.107) with the following settings: mismatch cost = 2; insertion cost = 3; deletion cost = 3; length fraction = 0.8; similarity fraction = 0.8. Sequencing raw data and detailed tables with expression values TPM, RPKM, total and unique gene reads for each sample are deposited under the GEO accession number GSE334666.

### Ligature-induced periodontitis

8–12 week-old mice were anesthetized and 5/0 silk sutures (SMI, Belgium) were looped and tied around both first maxillary molars using a triple knot as previously described (Abe and Hajishengallis 2013). Ligatures were assessed every other day and re-installed if absent. 7 days post ligature placement, mice were sacrificed, oral tissues were isolated for further analysis and maxillae analyzed using micro-computed tomography (μCT) for bone loss measurement.

### Micro-computed tomography (μCT) analysis

Maxillae were scanned using a high-resolution scanner (μCT 40, Scanco Medical AG, Bassersdorf, Switzerland). Measurements were taken at an operating voltage of 70kVp and 114μA current, with an exposure time of 200ms and voxel resolution of 12μm in all three spatial dimensions. To precisely quantify volumetric bone loss, quantitative three-dimensional measurements of teeth were performed. DICOM files were extracted from the scanner and analyzed in the OnDemand3D Application. The area between the cementoenamel junction (CEJ) and the alveolar bone crest (ABC) was marked and measured on the buccal and the palate sides.

## Acknowledgments

We thank Bettina Kalt for expert technical assistance.

## Abbreviations

LC: Langerhans cells
LN: lymph nodes
FACS: fluorescence activated cell sorting
LIP: Ligature-induced periodontitis

---

**Supplementary Figure 1:**
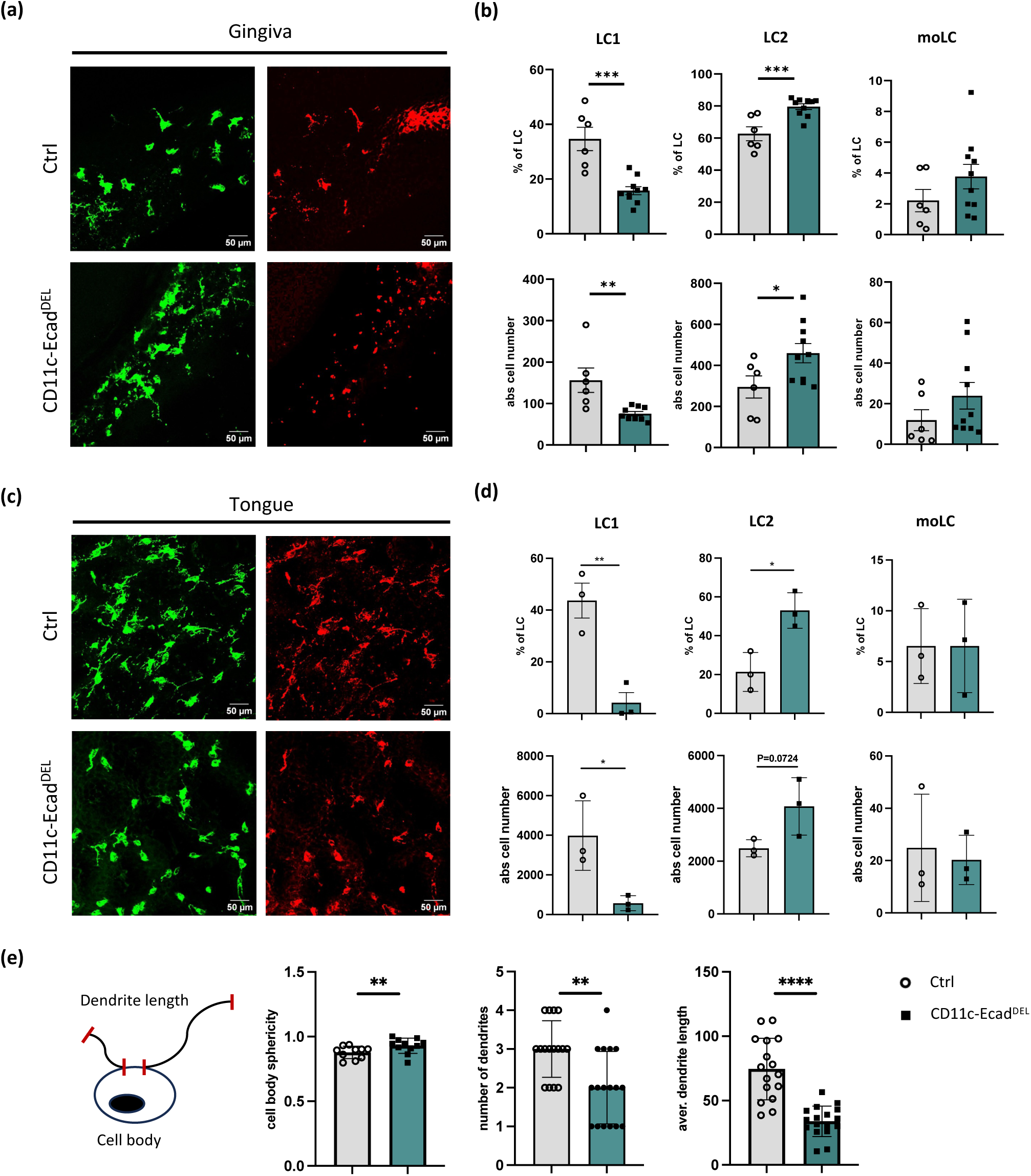
E-cadherin deficiency alters the morphology of LC throughout the oral mucosa. (a) Representative immunofluorescence images of epithelial sheets isolated from the gingiva of control and CD11c-Ecad^DEL^ mice, stained for MHCII (green) and Langerin (red). (b) Flow cytometric analysis and quantification of gingival LC subsets. (c) Representative immunofluorescence images of tongue epithelial sheets from control and CD11c-Ecad^DEL^ mice stained for MHCII (green) and Langerin (red). (d) Flow cytometric analysis and quantification of LC subsets in the tongue epithelium. (e) Quantification of LC morphology parameters in tongue epithelial sheets. Data are shown as mean ± SEM. Each symbol represents one mouse. Statistical significance was determined using unpaired two-tailed Student’s t-test or Mann–Whitney test as appropriate (* P < 0.05, ** P < 0.01, *** P < 0.001, **** P < 0.0001).

**Supplementary Figure 2:**
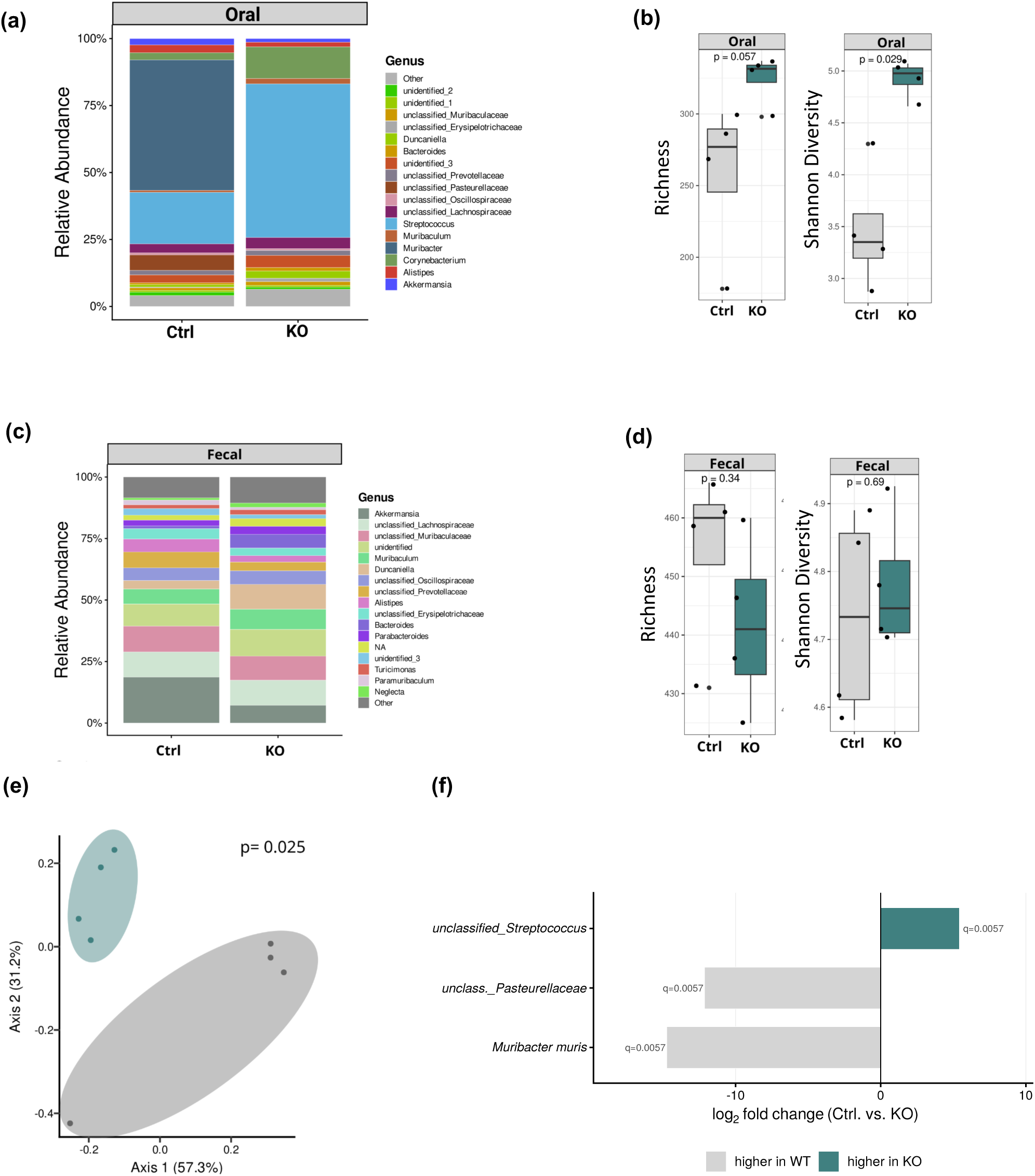
E-cadherin deficiency on LC induces oral dysbiosis. (a-f) Microbiota analysis of 32 week-old CD11c-Ecad^DEL^ and control mice. (a) Relative abundance of the oral microbiota at the genus-level, determined by 16S rRNA gene amplicon sequencing. (b) Oral alpha diversity (observed richness and Shannon diversity). (c) Relative abundance of the fecal microbiota at the genus-level. (d) Fecal alpha diversity (observed richness and Shannon diversity). Data are presented as boxplots (median, interquartile range, and individual mice), n = [4] per group. (e) Beta diversity of the oral microbiota, presented as principal coordinates analysis (PCoA) of generalized UniFrac distances; axis labels indicate the proportion of variance explained. (f) Differentially abundant genera in the oral microbiota, identified by LinDA, displayed as log2-fold change (control vs. CD11c-Ecad^DEL^); colored bars denote genera significant at padj < 0.05, with q-values indicated. Statistical significance: (b, d) Wilcoxon rank-sum test; (e) PERMANOVA; (f) LinDA with Benjamini–Hochberg correction. **p < 0.01; exact p-values are given where applicable.

**Supplementary Figure 3:**
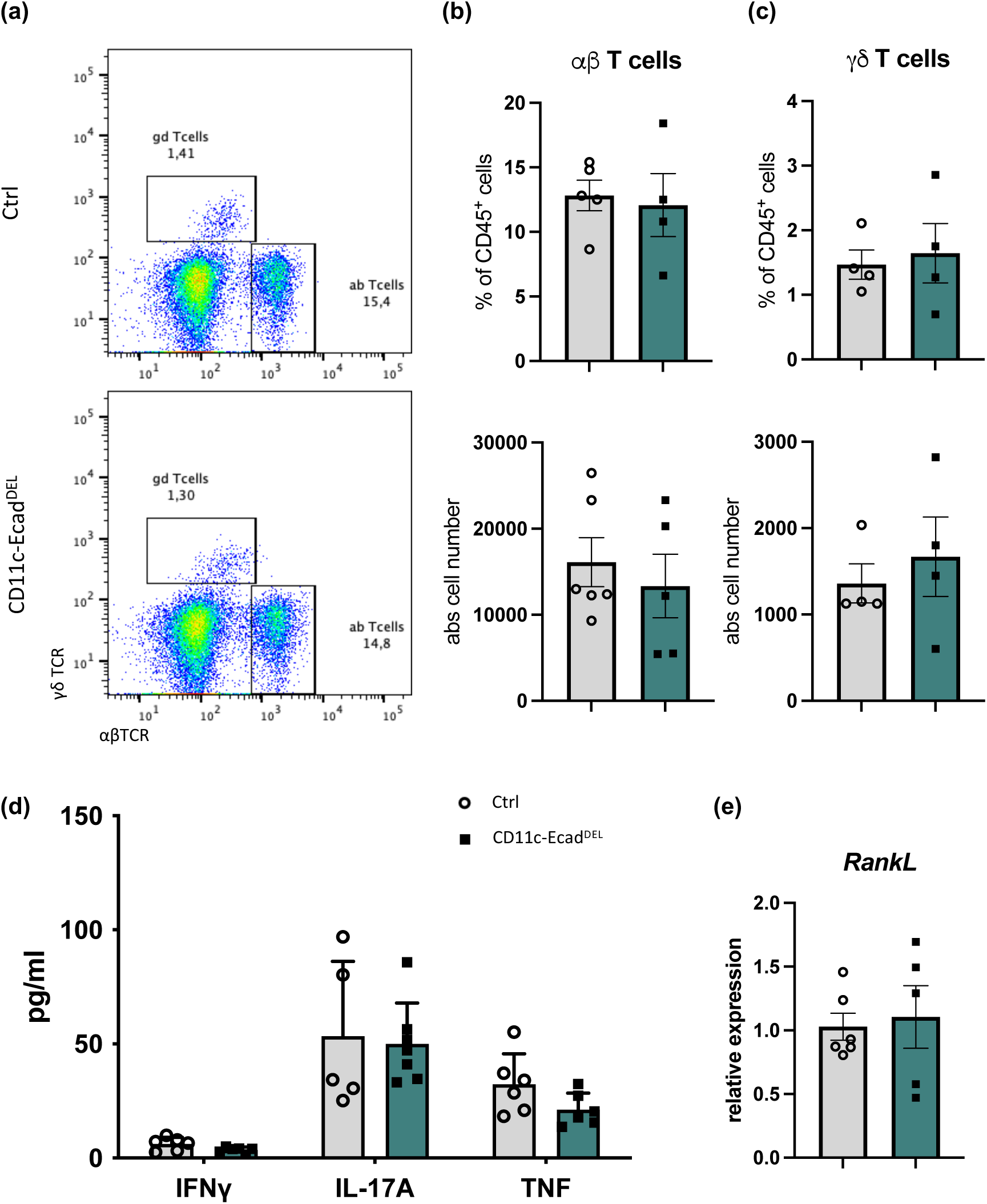
**E-cadherin deficiency in LC does not lead to inflammation and bone loss in young mice. (**a) Representative flow cytometry plots of gingival immune cells depicting αβ and ɣδ T cells of 12 week-old control and CD11c-Ecad^DEL^ mice. Quantification of (b) αβ T cells and (c) ɣδ T cells in the gingiva presented as frequencies and absolute cell numbers. (d) Gingiva IFNɣ, IL-17A and TNF cytokine levels as determined by Luminex multiplex assay. (e) Relative expression of *RankL* in the gingiva of 12 week-old control and CD11c-Ecad^DEL^ mice, measured by qRT-PCR.

